# Dual Computational Systems in the Development and Evolution of Mammalian Brains

**DOI:** 10.1101/2024.11.19.624321

**Authors:** Nabil Imam, Matthew Kielo, Brandon M. Trude, Barbara L. Finlay

## Abstract

Analyses of brain sizes across mammalian taxonomic groups reveal a consistent pattern of covariation between major brain components, including a robust inverse covariation between the limbic system and the neocortex. To find the functional basis of this inverse relationship, we mapped the multidimensional representations of task-optimized machine learning systems onto two-dimensional surfaces resembling the forebrain cortices. We found that visual, somatosensory and auditory systems develop ordered spatiotopic maps where units draw information from localized regions of the sensory input. Olfactory and relational memory systems, in contrast, develop fractured maps with distributed patterns of information convergence. Evolutionary optimization of multimodal systems for varying task objectives results in inverse covariation between spatiotopic and disordered system components that compete for representational space. These results suggest that the observed pattern of covariation between brain components reflect an essential computational duality in brain evolution.

---

One entity in the vertebrate brain has attracted the attention of researchers for decades, but it has resolutely resisted definition, or even exact specification of its contents. The anatomical term “limbic system” designates the presence of a number of the entity’s elements on the limbus, or edge of the embryonic lateral ventricle of the forebrain, including the rhinal and entorhinal cortices (variably, depending on each researcher’s focus), the subicular cortices and hippocampus, the amygdala, septal and other basal forebrain nuclei, with the connections to the cingulate cortices of the neocortex often emphasized. The “rhinencephalon”, smell-brain, features the olfactory system, the sensory receptors of main olfactory bulb and vomeronasal organ, and their primary targets, including primary olfactory cortex (“pyriform cortex”), the amygdala, additional basal forebrain nuclei of the olfactory tubercle and the parahippocampal gyrus, and the secondary targets of the hippocampus, thalamus and areas of neocortex as recruited. Those describing a memory-dedicated system will excise primary chemosensory and most basal forebrain nuclei from the group and add details and subdivisions to the hippocampus, entorhinal cortex and immediately interwoven cortex-like structures. Particularly in the field of neurology, the emotional sequelae of damage to collections of these regions were marked (e.g. “Kluver-Bucy syndrome”, “septal rage”).

Even with these disparate and often disjoint functions, integration over these nuclei and cortices was and is routinely emphasized. For example, “Papez’s circuit”, a collection of tracts often featured in medical textbooks, is so prominent it can be traced by blunt dissection, linking the hippocampus via the fornix to the hypothalamus, then to the anterior nucleus of the thalamus, to limbic cortices and back to hippocampus. It’s the rare current neuroscience textbook that fails to note the special emotional salience of olfactory memories. Unfortunately, one attempt to integrate over these diverse regions, Paul MacLean’s “triune brain” theory, popularized in Carl Sagan’s Dragons of Eden, persists to this day in the evocation of “my lizard brain” to account for uncontrolled emotional excess by the utterly false evolutionary tale that reptiles have only brain up to and including the olfactory and emotional rhinencephalon, grown over later by the logical neocortex, reaching its full flower in humans.

Fortunately, theories much better integrated over evolutionary, embryonic, neurophysiological and neurologic reality were developed. Walle Nauta, saw the whole brain as characterized by two ascending systems, one, the ascending core, open, polysynaptic, concerned with evaluative sensation and physiological maintenance, and that integrated at the level of the hypothalamus and limbic regions, and a second fast-conducting, analytical (as in rapid skin sensation, vision and hearing), organized at the level of the thalamus and neocortex. The brain becomes integrated over these two systems for movement selection and affect in the basal ganglia of the forebrain. A separate thread ranks cortical structures across both limbic and neocortical origins in a hierarchy of prediction and control of emotional/physiological, motor and sensory states, choices and evaluation. A characterization of the forebrain based on developmental gene expression gives a better original segmentation, and characterization of changing segmental and neuronal fates as brains enlarge. In fact, the need for any complementary characterization of forebrain at all is called into question. To list just a few of the contradictions in the limbic/other cortex distinction: no empirical justification exists to view olfaction as more emotion-laden than the voices of significant others; the hippocampus gets rather little olfactory input in many mammals and can’t be viewed as an olfactory structure; the hippocampus and neocortex both span the domains of emotion, event and sensorimotor world evaluation.

And yet, in a quite unexpected turn, the evolution and development of the vertebrate brain forces a two-systems view back into prominence. This is no invocation of the new brain-on-old brain layering of a scala naturae of fish, reptile, monkey and human – all vertebrates have essentially the same gross forebrain-to-spinal-cord segmentation, with no segments added on. Within those segments, the general functional commitment of regions remains remarkably stable, with the anterior-most and lateral-most regions of the embryonic neural plate destined to proliferate relatively the most in the larger brain. Rather, a strong pattern of complementary covariation between the cortical elements of the forebrain was first described quantitatively by Harry Jerison (though arguably misnamed) and has proven to be a volumetric distinction echoed between all large vertebrate taxonomic groups that have been studied, within taxonomic groups, and in individual variation within species, including our own. A push-pull relationship between the limbic system and the neocortex is evident in the data, with a volumetric expansion of one concomitant with a shrinkage of the other, after concerted allometric scaling is taken into account. We argue that this two-systems organization is characterized by distinct connectivity patterns in the component networks. The olfactory system and hippocampus have as their base a widely distributed patten of axon extension, whereby distributed neural activity patterns identify a particular odorant in the case of the olfactory structures, and a unique event in the case of the hippocampus. The neocortex, in contrast, preserves local spatial relationships in the projections of its subregions, cortical areas, onto each other. In most of the lateral convexity of the cortex, the dimensions that are mapped between areas represent egocentric space, over several sensory modalities and the motor modality, with cortical maps maintaining nearest neighbor relationships of the sensory and motor surfaces. In part of the cortex (the temporal lobe in humans), spatially ordered cortical projections are employed for non-egocentric mapping, as in tonotopy in auditory cortical representations, or the often-hybrid representations of faces, scenes, words and concepts.

We propose that the reason for the different patterns of axonal projections in the two systems is a computational one, that optimal representations of the associated modalities are achieved with distinct network architectures that induce system-specific inductive biases. To investigate, we encoded the representational space of task-optimized artificial neural networks in a two-dimensional model cortical sheet using topology-preserving maps. We found that networks optimized for visual, tactile and auditory representations converge to orderly spatiotopic maps that maintain nearest neighbor relationships of the input array, resembling sensory maps and connectivity profiles in the neocortex. In contrast, networks optimized for olfactory and relational memory representations evolved fractured maps with distributed connectivity profiles resembling axon distribution patterns of the olfactory-limbic system. Using evolutionary optimization, we optimized a multimodal network consisting of two subnetworks, one with spatiotopic and the other with distributed architecture. Selecting for different task objectives, we found an inverse relationship between the two subnetworks, resembling the inverse covariation between olfactory-limbic and neocortical system sizes observed

across species. Our results provide a computational account of a 550-million-year-old essential duality of the vertebrate brain.

## Neocortex and limbic system covariation

We used data collected for allometric analysis of brain structures in 183 mammalian species across 10 taxonomic groups including *Homo sapiens* [1–6]. Figure 1a plots the size (volume) of the neocortex against the size of the olfactory system for each species; the size of each structure is normalized with respect to core brain components (Methods) [7]. A robust negative correlation between the two systems is evident. Insectivores (hedgehogs, tree shrews, frogs) have relatively large olfactory systems and small neocortices, whereas primates, pinnipeds, ungulates and carnivores, and the manatee have reduced olfactory systems and enlarged neocortices. This inverse relationship can be partly understood by the dominant sensory systems of the taxonomic groups involved. Insectivores, or small insect-eating mammals, are often nocturnal, rely on olfactory cues for foraging, and have relatively large olfactory bulbs and olfactory cortices. In contrast, primates are highly visual, with a large fraction of the neocortex taken up by primary and extrastriate visual cortices. This contrast between olfaction and vision can be seen in all vertebrates, in aquatic species, reptiles, birds, and mammals, and is so pronounced that it can be seen in whole brains. Figure 1b shows a lateral view of the brains of an owl monkey and an agouti [2]. The brains are similar in volume, but the difference in the relative sizes of the neocortex and the olfactory system is apparent, with the neocortex in the primate obscuring both the olfactory system and the cerebellum from view.

**Figure 1:**
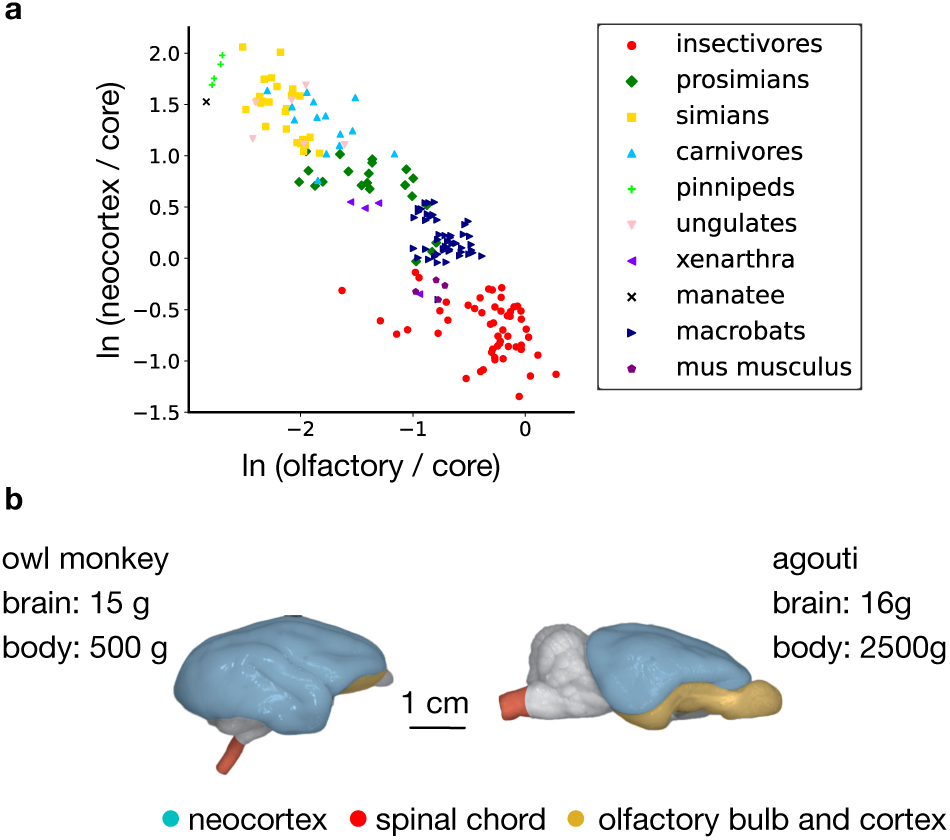
(a) Volume of the neocortex plotted against the volume of the olfactory system for 183 mammalian species across 10 taxonomic groups. The volumes are expressed as a fraction of the total volume of a core set of brain components (medulla, mesencephalon, diencephalon, striatum), in natural log scale. The olfactory system shown is the combined volume of olfactory bulb, olfactory cortex, olfactory tubercle including other medial forebrain nuclei, and the amygdala [1]. (b) Lateral view of the brains of an owl monkey (Aotus azarae, a small nocturnal primate) and an agouti (Dasiprocta agouti, a large diurnal/nocturnal rodent), chosen to demonstrate the strong contrast that can be seen in the relative size of the olfactory system versus the neocortex in brains of similar overall volume. Redrawn with permission from [2].

Figure 2a depicts a coronal section of the forebrain at the posterior thalamus level in a squirrel monkey (simian) and a nine-banded armadillo (xenarthra). The relatively larger olfactory cortices in the armadillo might be expected from their relatively greater reliance on olfaction, but its massive hippocampus is striking. In other mammals like the armadillo, olfactory input to the hippocampus is strong, but not dominating. The latter integrates multiple streams of information, including those originating in the neocortex, and thus is not a secondary “olfactory” structure [1]. Across taxonomic groups, the hippocampus positively covaries with the olfactory system (Figure 2b), which contrasts with the negative covariation between the olfactory system and the neocortex described above (Figure 1a). Why does the size of the hippocampus strongly covary with that of the olfactory system? In the following sections we argue that this covariation reflects a common computational architecture characterized by a common pattern of network connectivity distinct from that of the neocortex.

**Figure 2:**
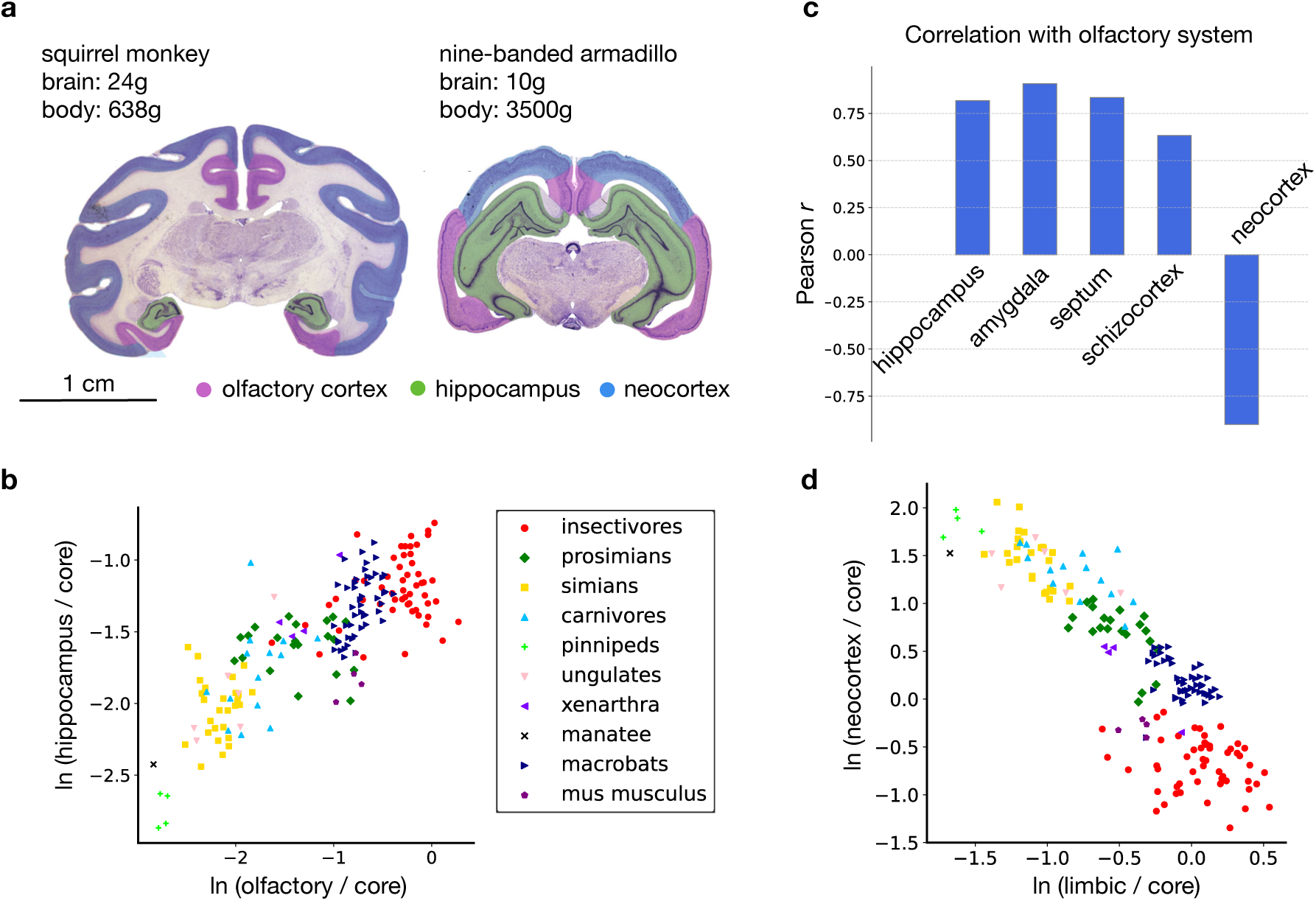
(a) Cross-section of the brain of a squirrel monkey (Saimiri sciureus) and a nine-banded armadillo (Dasipus novencinctus) showing the relatively enlarged neocortex in the monkey (blue) and the relatively enlarged olfactory and limbic cortex (dark purple) in the armadillo. Note also the relative expansion of the hippocampus in the armadillo (green). (b) Normalized volumes of the hippocampus plotted against that of the olfactory system across species (Pearson r = 0.82). (c) All limbic system components have strong positive covariation with the olfactory system. In contrast, the neocortex has strong negative covariation. Shown are the Pearson correlation coefficients between the normalized volumes of each structure and that of the olfactory system across species. “Schizocortex” includes the entorhinal, perirhinal, presubicular and parasubicular cortices. Amygdala volumes were available separately from the olfactory system for 103 of the 183 species analyzed. (d) Normalized volume of the neocortex plotted against that of the limbic system (sum of normalized volumes of all limbic system components) across species. A robust negative covariation is evident (Pearson r = -0.84). Images in subplot *a* are from the Comparative Mammalian Brain Collections, www.brainmuseum.org, property of the University of Wisconsin and Michigan State Comparative Mammalian Brain Collections, funded by the National Science Foundation and the National Institutes of Health.

In addition to the hippocampus, all other limbic system components, including the amygdala, septum, and schizocortex, positively covary with the olfactory system (Figure 2c) and with each other. This concerted change of limbic system components along with their inverse covariation with the neocortex (Figure 2d) suggests a two-systems view of the macroscale organization of the brain. In early neurodevelopment, the two regions can be easily distinguished by their patterns of cell proliferation and early gene expression [8] and in the expression of transcription factors, notably LAMP, “limbic-associated marker protein” [9], suggesting distinct developmental rules of network formation.

## Structure of sensory and cognitive maps

### Information structure and optimal representations

To understand the reasons underlying the pattern of covariation in brain evolution described above, we investigate the structure of distinct information sources driving brain computation and their optimal representations. For this purpose, we assess the multidimensional representational space of modern machine learning systems, optimized for one of four sensory modalities – vision, audition, somatosensation and olfaction – followed by a system optimized for relational memory. The sensory systems consist of the following artificial neural network architectures pre-trained on large data corpora **(1)** ResNet trained on images from the ImageNet database [10, 11], **(2)** wav2vec 2.0 trained on natural speech from the Libri speech corpus [12, 13], **(3)** a convolutional architecture trained on a dataset of tactile signals of the human grasp [14], and **(4)** an olfactory systems model trained on olfactory receptor neuron activation profiles [15]. These networks show strong correspondence to neural activity and structural connectivity patterns in the brain within particular parameter regimes [11, 13, 15]. Training optimizes network connection weights, and subsequently, the networks embed manifolds of stimulus feature information in their activity.

The data used to optimize each model is depicted in Figures 3a-d. We observe a dichotomy in the structure of information across modalities, with visual, auditory, and somatosensory data showing pronounced spatial structure, as measured by spatial autocorrelation (Moran’s I [16]), and olfactory data lacking any such structure (Figure 3e). Coarse-scale chemotopy in olfactory receptor neuron population representations have been proposed in some studies [17], but it is clear that they lack the fine-grained spatial order characteristic of primary visual, somatosensory and auditory representations [18–21]. We argue that this dichotomy in the structure of information sources underlies the two-systems architecture that we proposed, each system pre-configured with knowledge of the core statistical structure of their input.

**Figure 3:**
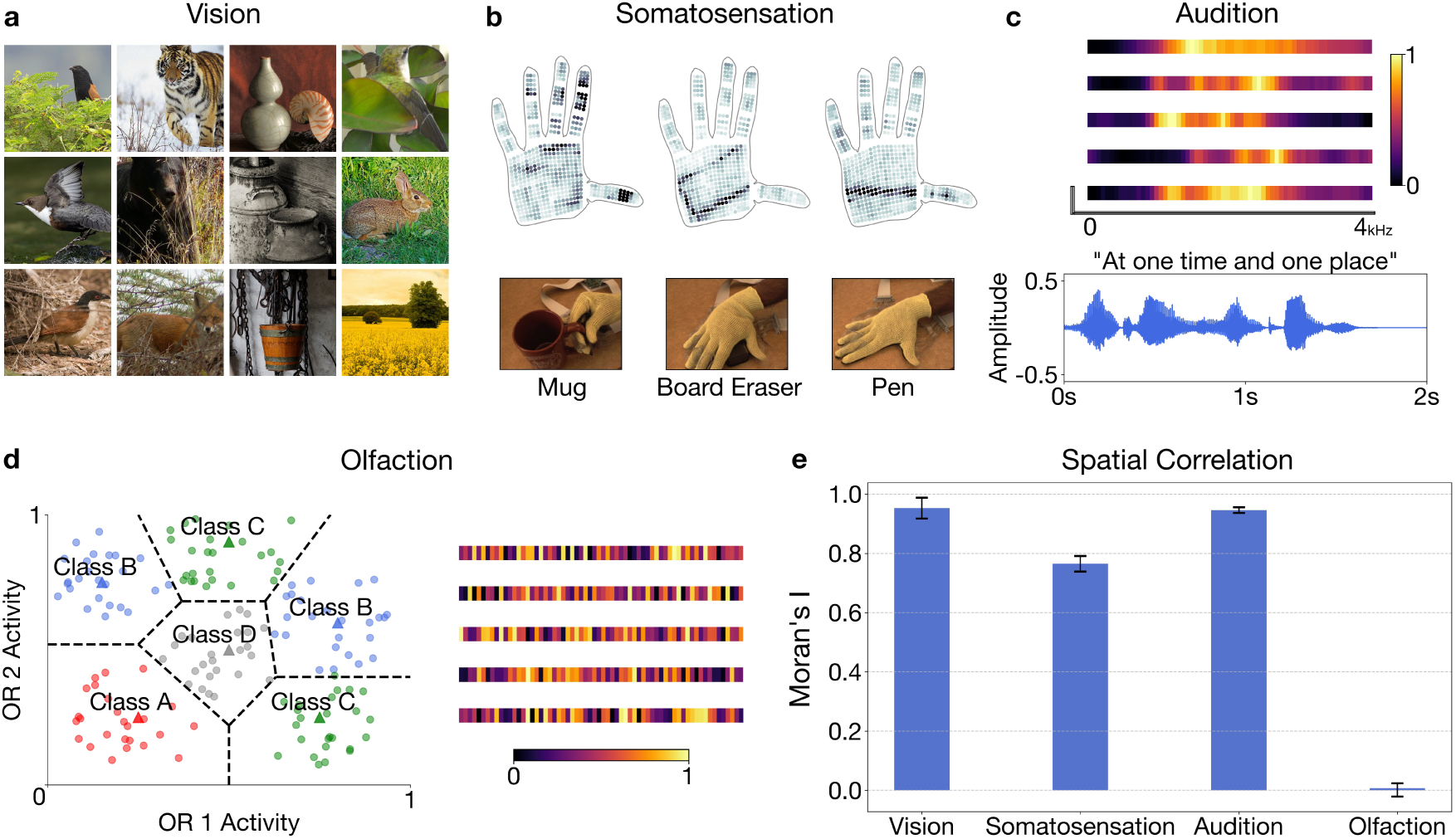
Information sources of four sensory modalities. (a) Samples from the ImageNet dataset. ImageNet images are characterized by spatial correlation visible in the form of contrastive backgrounds, repeating textures, and distinct natural shapes. (b) Top: visualizations of pressure readouts from the tactile glove dataset [14]. Bottom: perspective images showing the creation of each sample. Activated regions of the tactile glove are highly localized, as clusters of adjacent sensors respond to pointwise contact with the object being classified. (c) One-dimensional cochleagram slices (top) and time-amplitude representation of a single sample (bottom), each from the LibriSpeech ASR corpus. In the cochleagram slices, frequency bands associated with human speech form localized patterns. (d) Left: subset of 150 synthetic odors from the Wang et al dataset [15] showing activation values of two olfactory receptor neurons (ORNs). Each odor sample is encoded by 50 ORNs in the dataset. Individual odors are classified by their proximity to two prototype odors (triangles) which are distributed across the full range of possible values and exhibit minimal local spatial correlations. Right: ORN activation profiles of five sample odors from the dataset. (e) Barplot showing spatial autocorrelation (Moran’s I) across the aforementioned datasets. Moran’s I is the ratio of the spatially weighted covariance to the overall variance, normalized by the number of observations and the sum of the spatial weights. This metric is expressed in a range from -1 to 1, where positive values indicate increasingly strong local similarity, values near zero suggest random spatial patterns, and negative values indicate strong local dissimilarity.

### Spatiotopic and disordered maps

After optimization of network parameters in the models, network activity constitutes an optimized population code of stimulus features across a multi-dimensional feature space. A prevailing theoretical account of cortical organization posit that such multi-dimensional feature spaces are represented in two-dimensional cortical surfaces in a manner that preserves the topology of the stimulus manifold [22–25]. Similar stimulus features are represented in nearby locations on the cortical surface to aid efficient similarity-dependent computations. Following this account, we construct Kohonen self-organizing maps (SOMs) to represent the optimized feature spaces of our models on a two-dimensional cortex-like surface [25]. We consider the organization of primary sensory maps that are in place early in brain development and whose activity drive areal organization across the system [26, 27]. Specific layers of our models show strongest correspondence to activity in primary sensory areas and we construct SOMs using activity in these layers (Methods) [11, 13–15].

Figure 4a shows the maps that develop in each modality, highlighting two contrasting structures. Visual, auditory and somatosensory maps converge to orderly representations that maintain nearest neighbor relationships of the sensory surface resembling spatiotopic organization that characterizes the entire neocortex. Olfactory maps, in contrast, develop fractured representations akin to disordered chemotopy in the olfactory bulb and cortex. We quantify the degree of global spatial order in each map by computing the mean Euclidean distance between each unit’s preferred stimulus and that of its adjacent units, normalized by the expected distance in a random arrangement and negated to result in an order metric (Methods). This results in an index that ranges from 0 (low order) to 1 (high order). We measure values of 0.90, 0.86 and 0.89 for visual, somatosensory and auditory maps respectively, and 0.11 for olfactory maps.

**Figure 4:**
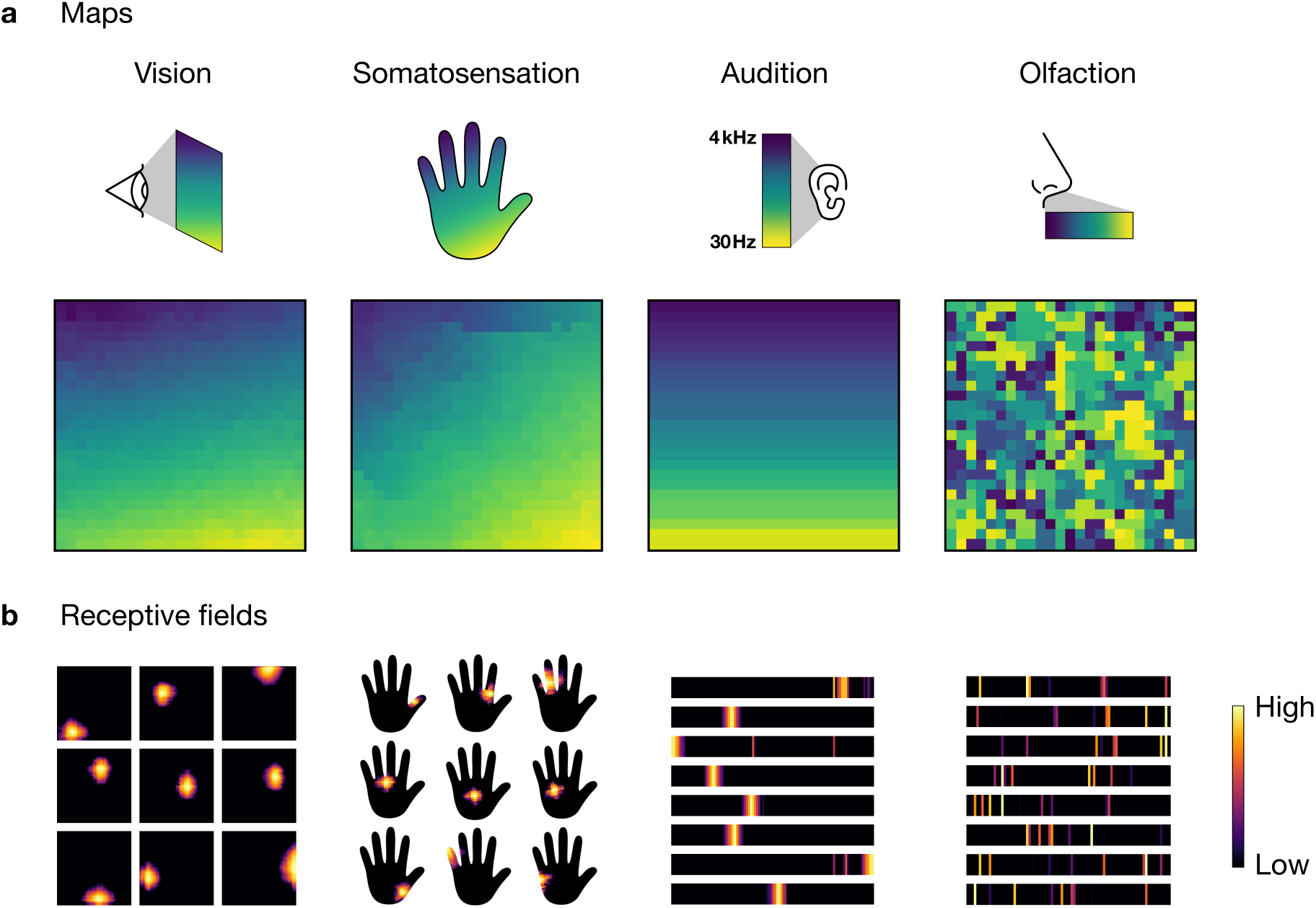
(a) Model cortical sheets constructed by encoding the multidimensional representational space of task-optimized artificial neural networks in two-dimensional surfaces using Kohonen self-organizing maps [25]. Models of four sensory systems are constructed. The top row depicts the sensory input colored by location. The bottom row shows maps of the sensory input on the model sheets. Each sheet consists of 25 x 25 units. Each unit is colored according to the input location that maximally activates the unit. The gradual transition of colors in visual, somatosensory, and auditory representations indicate a smooth mapping of the input space on the cortical surface. No such mappings are found for olfactory representations. (b) Example receptive fields of model units indicating the degree to which each region of the input activates a unit. Receptive fields in vision, somatosensation and audition are spatially localized whereas those in olfaction are spatially distributed.

We then assess the receptive fields of individual SOM units and find that visual, somatosensory and auditory units draw information from localized regions of the sensory surface, whereas olfactory units sample across distributed parts (Figure 4b), recapitulating a central distinction between afferent projection patterns in primary olfactory cortex and primary sensory areas of neocortex. We measure the degree of locality of the receptive fields by computing the spatial autocorrelation (global Moran’s I) of representative SOM unit receptive fields, again resulting in an index that ranges from 0 (low locality) to 1(high locality). We calculate locality indices of 0.97, 0.84 and 0.99 for vision, somatosensation and audition respectively, and 0.00 for olfaction. Further, we find that this pattern of information convergence arises within unstructured (fully-connected) networks solely as a consequence of gradient-based optimization of information representation without prespecified network structure (Supplementary Materials) [28]. Overall, the contrasting structure of information representation between the spatiotopic and olfactory modalities mirrors the dual structure of their information sources (Figure 3) and underscores a pronounced duality in the organization of optimal sensory representations.

### Relational memory

We next assess representations in the Tolman-Eichenbaum Machine (TEM), an artificial neural network for relational inferences modeled on computations in the hippocampus and entorhinal cortex [29]. The network is trained to learn conjunctions of perceptual and structural representations and develops a cognitive map via which relationships between objects and their spatial or abstract locations are inferred (Figure 5a). Activity within the network resemble activity patterns in spatial and non-spatial cell types in the hippocampus and entorhinal cortex, as well as their structural remapping under new stimuli, within particular parameter regimes [29]. We encoded the conjunctive representations of this model in a two-dimensional surface using a SOM and compared its structural features to the sensory systems maps of Figure 4.

**Figure 5:**
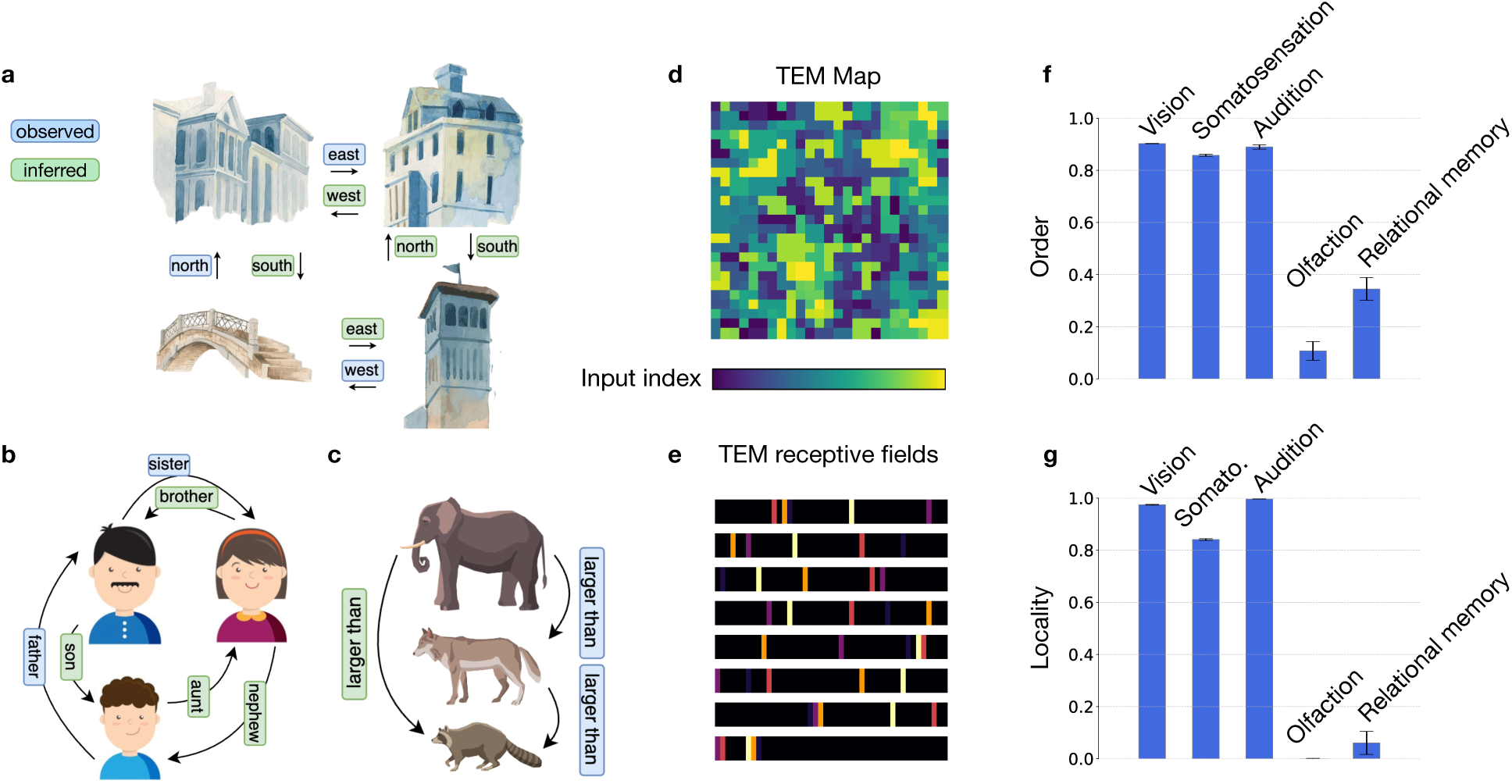
The Tolman-Eichenbaum Machine (TEM) learns relational structure of a problem and uses that knowledge to make quick inferences in new problem instances. Relational structure exists in many situations, for example (a) spatial reasoning, (b) social hierarchies and (c) transitive inference. The TEM infers the green relationships after observing only the blue ones using a generative model of the hippocampus and entorhinal cortex. Figures adapted from [29]. Clipart designed by Freepik. (d) Hippocampal activity of the TEM is encoded in a two-dimensional surface using Kohonen self-organizing maps [25]. Shown is a surface consisting of 25×25 units, where each unit is colored according to the location of the input that maximally activates that unit. The disordered map resembles the olfactory map of Figure 4a. (e) Example receptive fields of hippocampal units in the TEM. The receptive field of a unit (1D horizontal bar) shows the degree to which each region of the input activates that unit. The spatially dispersed receptive fields resemble those of the olfactory map in Figure 4b. (f-g) Indices measuring the order of the maps and the locality of the receptive fields across the five modalities. Hippocampal relational memory and olfactory representations have distinct organization compared to the spatiotopic sensory systems as indicated by the values of these indices. Plotted are the means and standard deviations across ten simulations with different random seeds.

As in the olfactory maps, hippocampal maps develop fractured representations that lack global spatial order, with SOM units drawing information from distributed parts of the TEM’s input (Figure 5d-e), resembling sparse and distributed coding in the hippocampus [30–32]. We compare map indices across models and observe that both olfactory and hippocampal representations exhibit low spatiotopic order and distributed receptive fields, indicating that the associated computations benefit from inductive biases that are distinct from those of the other sensory systems (Figure 5f-g).

## Evolution of a dual computational system

The structure of optimal representations discussed so far mirrors the structure of primary sensory representations in the brain in place early in development. This pre-configuration of network architectures prior to stimulus exposure serves subsequent computations by encoding knowledge of the structure of expected stimuli in network connectivity. What developmental factors shape this connectivity and what axes of variations are offered up to natural selection? To investigate, we configured a dual-systems architecture consisting of two subnetworks, one with spatiotopic and the other with distributed organization drawing input from five sources of information – visual, somatosensory and auditory sources providing input to the spatiotopic network, and olfactory and hippocampal sources to the distributed network (Figure 6a). Units within the spatiotopic and distributed networks are allocated equally among their constituent components. We ask how these networks may co-vary under selection pressures along a single dimension of variation – the location of the boundary dividing the spatiotopic and distributed subnetworks. The boundary determines the number of units, or the amount of computational resources, allocated to each modality and its variation accounts for a minimal degree of stable developmental control of tissue allocation (Figure 6b). We use an evolutionary algorithm [33, 34] to optimize the position of this boundary under a selection criterion defined by a parameter *s* that takes a value between 0 and 1; a low value of *s* selects for olfaction and a high value selects for vision. We initialize a population of networks, each with a roughly even allocation of units between the spatiotopic and distributed networks, and ran the evolutionary algorithm for fifty optimization steps. At each step, the networks with the highest task performance are selected, and the locations of their boundaries are varied uniformly at random to generate networks for the subsequent step (Methods). Task performance is a combination of olfactory and visual signal identification, as defined by the parameter *s*. The algorithm optimizes the performance of the networks across iterative steps of variation and selection.

**Figure 6:**
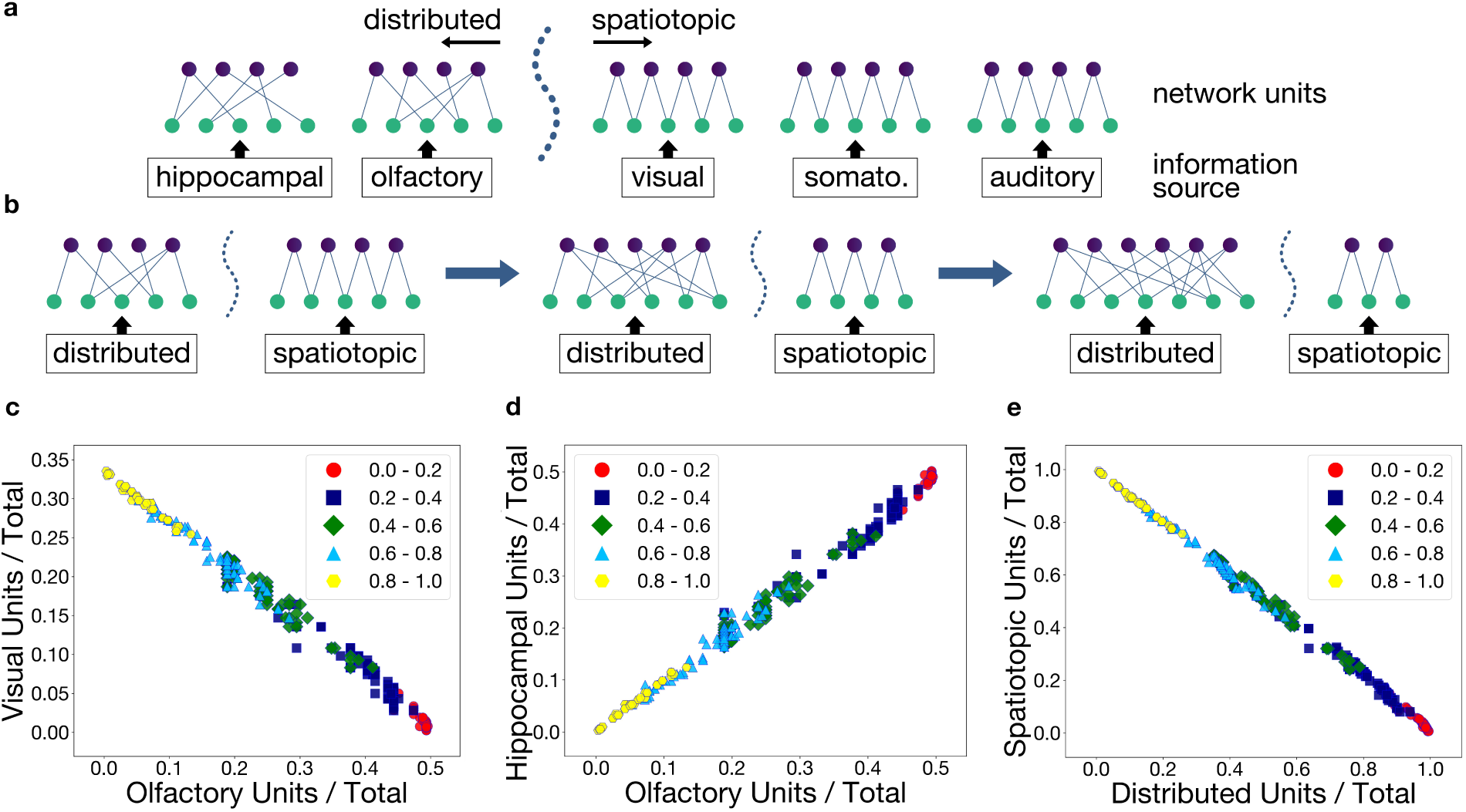
An evolutionary algorithm accounts for patterns of covariation among brain components. (a) A dotted boundary separates two networks with distributed connections from three networks with spatiotopic (local) connections at their inputs. Shifting this boundary adjusts the number of units allocated to each side proportionally. New generations of networks are produced by randomly stepping the boundary in the direction associated with the fittest survivors from the previous generation. (b) Depiction of selecting for olfaction. The boundary shifts in a direction that expands the distributed network resulting in a larger allocation of network units to the constituent components. (c) Allocation percentages to the visual and olfactory components after 50 iterations of variation and selection. Points are categorized by groupings of “*s*” values in a range from 0 to 1 where *s* is the weight assigned to the test performance of vision when calculating the fitness of a given network. (d) Similar to subplot *c*, but showing the allocation percentages between the hippocampal and olfactory components of the network.(e) Displays the combined unit allocations for the visual, somatosensory, and auditory portions of the network (y-axis) against the combined allocations for the olfactory and hippocampal portions (x-axis) after 50 iterations.

Figure 6c shows the outcome of the evolutionary optimization process for different values of *s*. When olfactory performance is selected for (low values of *s*), the number of units allocated to olfaction increases at the expense of vision, and vice-versa (high values of *s*), recounting the olfaction vs vision account of sensory specialization (Figure 1). We observe that instead of singular expansion of the selected component, a concerted change in the size of all components takes place, with olfactory and hippocampal network sizes positively covarying with one another and inversely covarying with the sizes of the spatiotopic networks (Figure 6d-e). These results show that a minimal degree of developmental control is sufficient to account for the overall patterns of covariation observed across species (Figures 1 and 2). A segmental ‘prosomeric’ structure of the forebrain defined by domains of regulatory gene expressions suggest a mechanism of this nature [35, 36].

When evolution selects for brain components, it selects for function that arises from underlying computational processes. Particular computations are aided by specific features of network architectures and representational schemes, the spatiotopic and distributed networks discussed here being examples from what may be an abundant space of architectural priors exploited in brain computation. Variations in network organization offered up to natural selection are shaped by developmental processes and thus constrained by directions of variation under developmental control. These ‘units of development’ might not precisely correspond to the function that is selected for, and the outcome of selection is likely a result of specific functional demands and available developmental variation, mediated strongly by metabolic costs and other contingencies [1]. The results presented here suggest that the pattern of covariation between the limbic system and the neocortex observed across species is a consequence of developmental control of tissue allocation between two computational systems offered as loci of selection. Despite the diversity of limbic system function, a common computational architecture and coarse developmental structure underlies concerted variation of its components. Selecting for one causes the selection of all, with pleiotropic effects on behavior.

## Acknowledgments

This work was supported by NSF grants 2223811 and 2319060 to NI.

## Methods

### Neuroanatomical data

Brain volume measurements of insectivores, prosimians and primates are taken from [4, 5], macrobats from [6]; carnivores, pinnipeds, ungulates, xenarthra, and the manatee from [1], and mus musculus from [3]. Together, these constitute an extensive dataset covering a wide range of mammalian species, a diverse array of niches (ranging from burrowing to flying, nocturnal and diurnal, omnivores and specialists), and a wide spectrum of brain sizes. Volumes are listed for a core set of brain components – medulla, mesencephalon, diencephalon, and striatum – as well as limbic system components, neocortex and cerebellum. Limbic system components are amygdala, olfactory system, septum, hippocampus, and entorhinal cortex and associated cortical structures (parahippocampal, presubicular, parasubicular and subicular cortices). Structure nomenclature and phylogenetic assignments are matched to the initial work of Stephan and colleagues [4, 5].

In Figures 1 and 2 of the main text, neocortex, olfactory system, and limbic system volumes are expressed as a fraction of the total volume of core brain components (medulla, mesencephalon, diencephalon, and striatum) [7]. Following Stephan [5] and Reep [1], the olfactory system is the combined volume of olfactory bulb, olfactory cortex, olfactory tubercle including other medial forebrain nuclei, and the amygdala. See Reep [1] for the minimal effects of inclusion or exclusion of the amygdala. Figure 1c (amygdala bar) considers the amygdala separately from the olfactory system in the 103 species for which the volumes are separately available.

The patterns of covariations in Figures 1 and 2 are also observed when considering deviations from allometric expectations (Figure S1 in Supplementary Materials).

### Self-organizing maps

Kohonen self-organizing maps (SOMs) are constructed from the multidimensional feature spaces of pre-trained deep neural networks [25]. The spatial organization of maps arising from five distinct networks are assessed, as described in the main text. The constructions of the maps are consistent across all the networks.

To construct each map, the corresponding network’s input space is divided into non-overlapping patches of static, with shapes and values scaled according to the size of the input. The set of patches are passed through the network, one at a time, and activations arising from the final feature extraction layer (specified below) are recorded. This constitutes the set of activations that are used for constructing the SOM. This will be referred to as set *A* below.

The SOM is initialized with 625 units arranged in a 25 x 25 square lattice. Elements from set *A* are then presented to the SOM as input, one at a time. Each SOM unit computes a weighted sum of this input; the weights are chosen uniformly at random at initialization. The most active SOM unit as well as units within its local neighborhood are then selected and their weights updated according to Kohonen’s algorithm [23]. The algorithm updates the weights such that nearby units develop similar weights over the course of multiple iterations through *A*. This process is repeated for a total of 100,000 iterations. The full algorithm is provided in Algorithm 1 below. A hyperparameter sweep is provided in Supplementary Materials (Figures S3-S4).

To visualize the map, a smooth color-gradient is set up to index all the patch locations that were used to construct *A*. After constructing the map using the algorithm described above, elements of *A* are again passed to the map, one at a time. The map unit that is most active for each element is assigned the corresponding patch’s color. The map is then visualized as a 25 × 25 heatmap, where each cell corresponds to a SOM unit and colored accordingly (Figure 4a of the main text).

The location preference of a map unit is defined as the patch index for which that unit had its highest activation. The spatial order (smoothness) for each map (Figure 5f of the main text) is quantified by examining the location preference of each map unit. For each unit, the average distance between its location preference and the location preferences of immediately adjacent units is calculated. This is done for all units in the map and averaged. This average distance is then divided by a normalization factor, representing the expected distance in a random map computed with 10,000 Monte Carlo samples. The resulting metric is then negated to quantify smoothness and range normalized to 0-1.

#### Algorithm 1 Self Organizing Map Algorithm

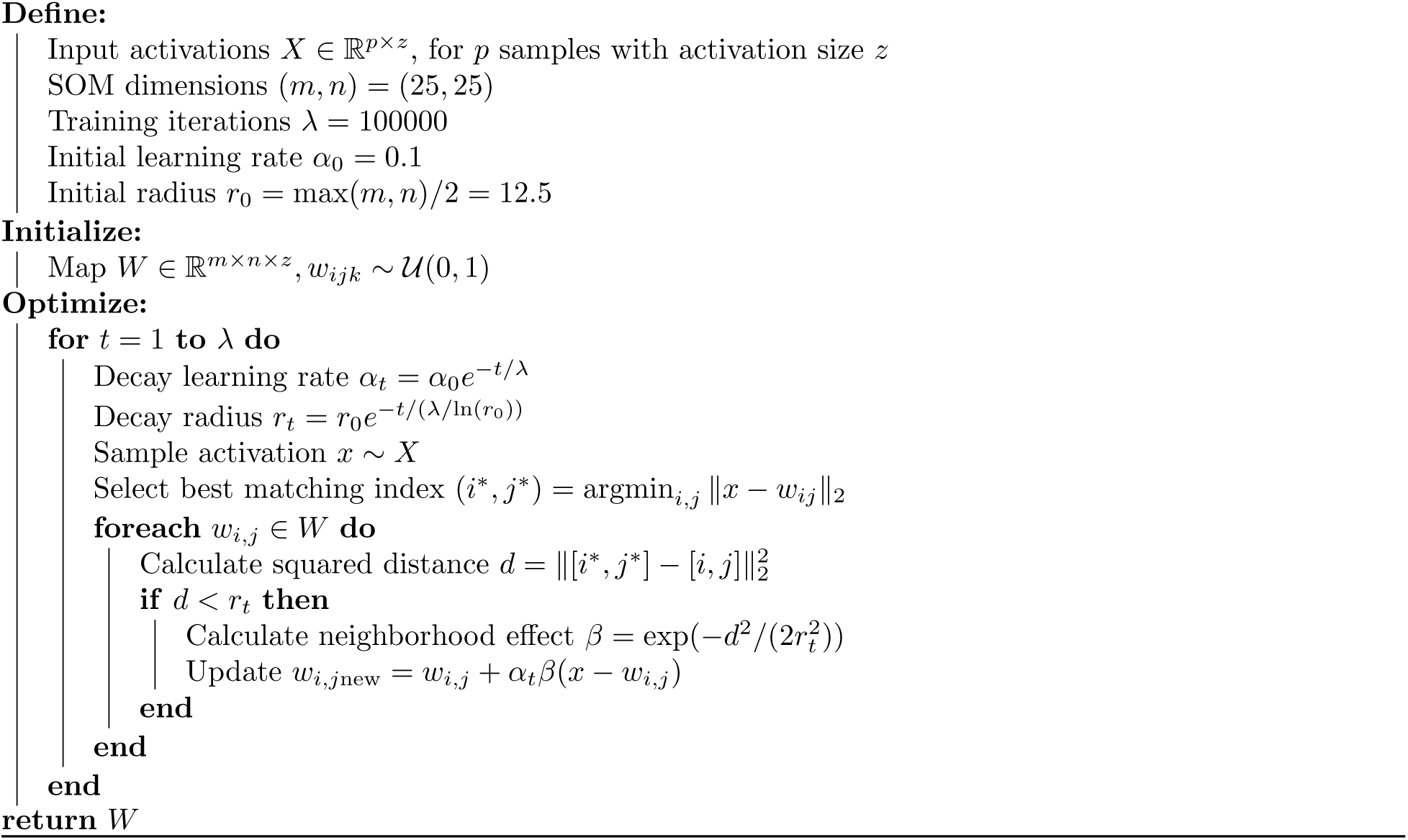

The receptive field of a map unit is visualized as a heatmap whose dimensions match the shape of the input data (Figure 4b of the main text). The intensity of each cell in the heatmap corresponds to the degree of activation of the map unit in response to input at that cell location. The top decile of activations are shown. The receptive field locality measure (Figure 5g of the main text) is calculated by averaging the spatial autocorrelation (global Moran’s I [16]) of the receptive fields of all units in the map.

The specific considerations made for each map are specified below.

The vision map is constructed using activity from a pretrained ResNet18 model [10, 11]. The activations are recorded from layer 4, after the final CNN feature extraction block. The raw data input size is 225 × 225 × 3. The patch size is 3×3×3, with patch elements assigned values between 0.5 to 1, chosen uniformly at random, resulting in 5,625 patches for constructing the map. Before presenting the dataset of patches to the network, the dataset is normalized to have the same mean as a subset of the ResNet18 training dataset.

The somatosensory map is constructed using a 1-frame grasp classification network, trained following the procedure outlined in [14]. Activations are recorded from layer 10, after the ResNet feature extraction block, maintaining consistency with the vision network. The raw input data shape is 32 × 32 × 1. The patch size is 1 × 1, with values between 0.5 and 1, chosen uniformly at random, resulting in 1,024 patches for constructing the map. The patches are filtered to only include patch coordinates with non-zero pressure values in the network’s training dataset.

The audition map is constructed using a pretrained wav2vec 2.0 model [12,13]. Activations are recorded from layer 7, after the feature projection layer of the encoder block, consistent with the somatosensory and vision networks. The raw waveform is truncated to 3,000 frames and transformed to a spectrogram with 5,000 frequency channels. Patches are created in the frequency domain, where each patch is a single frequency channel. Patch values are drawn uniformly at random from the range [0, 1], and then mapped to the range [11, 22], matching the distribution of the raw data. After patch construction, the data is transformed back to a waveform format. The SOM weight-update function is modified to accommodate the 1-dimensional frequency data by only considering a single dimension distance in the update function.

The olfactory map is constructed using the olfactory systems model in [15]. The input dimension of the model is increased from 50 to 4,096 in order to use the same SOM size as those in the other modalities. Activations are recorded after the third layer, corresponding to the pyriform cortex in the model. The patch size is 1 × 1, with values assigned between 0.5 to 1, uniformly at random. Before presenting the dataset of patches to the network, the dataset is normalized to have the same mean as the network’s training dataset.

The hippocampal map is constructed using activity in the Tolman-Eichenbaum Machine [29] with an observation dimension of 45. The network is probed using a single batch of 100 rollouts and 45 basis vectors (vectors of zeros with a single 1). At each step in the rollout, each basis vector is passed to the model and the activation of the 7th inference layer, corresponding to the calculation of the inferred grounded location, is recorded.

### Evolutionary algorithm

The evolutionary algorithm operates on the parameter allocations of a neural network architecture consisting of two subnetworks, one with spatiotopic and the other with distributed connections. Each subnetwork consists of multiple components – visual, auditory and somatosensory components in the spatiotopic network, and olfactory and hippocampal components in the distributed network. Each component has its own input, hidden, and output layers. The components are trained simultaneously in PyTorch using modality-specific datasets. Due to the computational requirements of evolutionary algorithms, smaller datasets are used for this assessment. These are CIFAR-10 for vision, Speech Commands for audition, and an associative memory dataset for the hippocampal subnetwork [32]. The somatosensory and olfactory datasets are the same as those in the other assessments.

In the setup described above, the spatiotopic network components feature input layers that draw information from a localized 5 × 5 region of the input, while the distributed network components have fully connected input layers with a random subset of weights set to zero. The size of this random subset is chosen such that the active parameters in the network roughly equal that of a locally connected network of the same size. Hidden layer sizes in the spatiotopic and distributed networks are determined by a boundary *B* between the two, whose position (indicated by the dotted line in Figure 6 of the main text) determines the allocation of network units. Units allocated to each subnetwork are distributed evenly among its components.

The evolutionary algorithm is run for fifty iterations, each with a population of fifty networks. The networks differed in the position of the boundary *B*, and thus in the allocation of units between the two subnetworks. First, a seed network is constructed in which the boundary is placed at a location that evenly divides the total available parameters across all five subnetworks. The initial population is constructed by sampling uniformly within ±10% of the seed network’s boundary location, with the seed network included by default in the first generation.

Next, the networks are trained with gradient descent for fifty epochs. The fitness of each network is measured using a weighted combination of task performance in the visual and olfactory modalities, described below. Twenty-five networks (half of the population) that have the highest fitness scores are then selected for the next generation, along with an additional twenty-five networks sampled with boundary values centered around that of the most fit network in the current generation. The results do not depend on particular choices of these settings; the same overall results are observed for a wide range of settings (Figure S5-S7 in Supplementary Materials).

The fitness function weighs task performance using a parameter *s*, where a value of 1 fully emphasizes visual performance and a value of 0 fully emphasizes olfactory performance. Varying this parameter between 0 and 1 produces the plots in Figure 6c-e of the main text, which show the top ten highest-performing networks after completion of the algorithm, given a specific value of the parameter *s*.

## Supplementary Materials

### Allometric expectation

Figure S1a plots the volume of the limbic system and the volume of the neocortex against the volume of core brain components (medulla, mesencephalon, diencephalon, and striatum) across 183 mammalian species. Both structures scale predictably at characteristic rates. The regression line running through each set of scatter points provides the allometrically expected volume of each structure, given the volume of core brain components. Deviations from this *allometric expectation* is measured by calculating the residuals of the scatter points with respect to their corresponding regression line. These residuals are shown in Figure S1b, which plots the residuals of the neocortex against those of the limbic system. As in Figure 2d of the main text, a negative covariation between the two structures is evident. Species with larger than expected neocortices have smaller than expected limbic systems (low limbic / high neocortex quadrant), whereas those with larger than expected limbic systems have smaller than expected neocortices (high limbic / low neocortex quadrant). One exception to this dominant trend is microbats (not shown), which have reductions in both components due to selection pressures for light-weight nimble flight [1].

**Figure S1:**
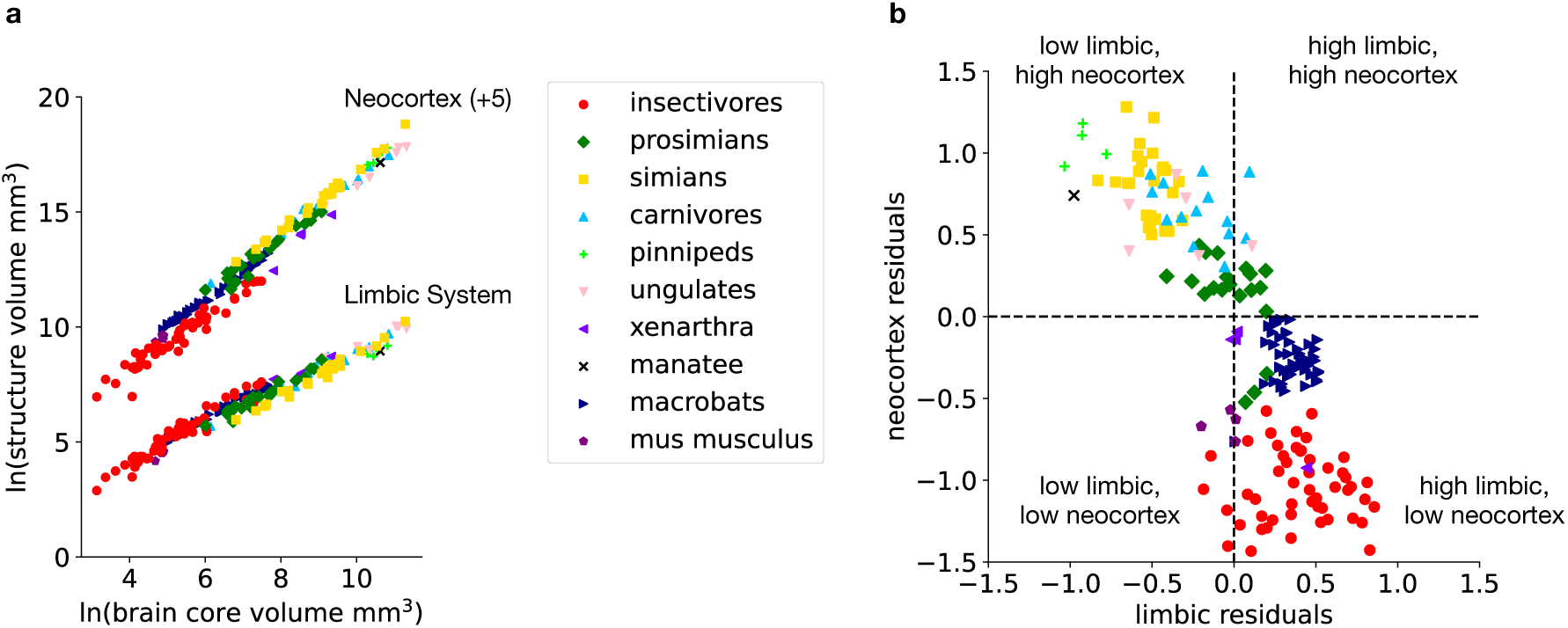
(a) Volumes of the neocortex and the limbic system plotted against the volume of core brain components across 183 species. An arbitrary constant of +5 is added to the neocortex data points to visually separate the plots. (b) Residuals of the neocortex and limbic system data points in subplot *a* inversely covary with each other.

### Optimizing unstructured networks

To investigate the patterns of information convergence arising in artificial neural networks that lack pre-specified network structure, three-layer fully connected networks are trained using the same datasets used in the evolutionary algorithm (Methods). Each network is trained with the “*β*-LASSO” objective described in [28], which appends a weight thresholding step to stochastic gradient descent. This approach encourages network weights to trend aggressively towards zero during training, creating networks that are highly sparse, and which approximately match the performance of identical networks trained with stochastic gradient descent while using only a fraction of the active (non-zero) parameters.

When evaluating these networks during training, we measure the level of sparsity in the first layer and tune the thresholding hyperparameter in *β*-LASSO to achieve a comparable number of active weights for each network. At the highest levels of sparsity (where 95-99% of weights are zero), the networks develop modality-specific patterns of connectivity with varying degrees of locality, reflecting the biases inherent in their respective datasets. Figure S2 shows the distribution of receptive field sizes in each modality after tuning for this objective. Networks trained on data with a high degree of local correlation (vision, somatosensation and audition) generally have localized connectivity consisting of small clusters of non-zero weights, while those trained on data without locally correlated structure learn distributed patterns of connectivity across the full span of their inputs (olfaction and hippocampus).

**Figure S2:**
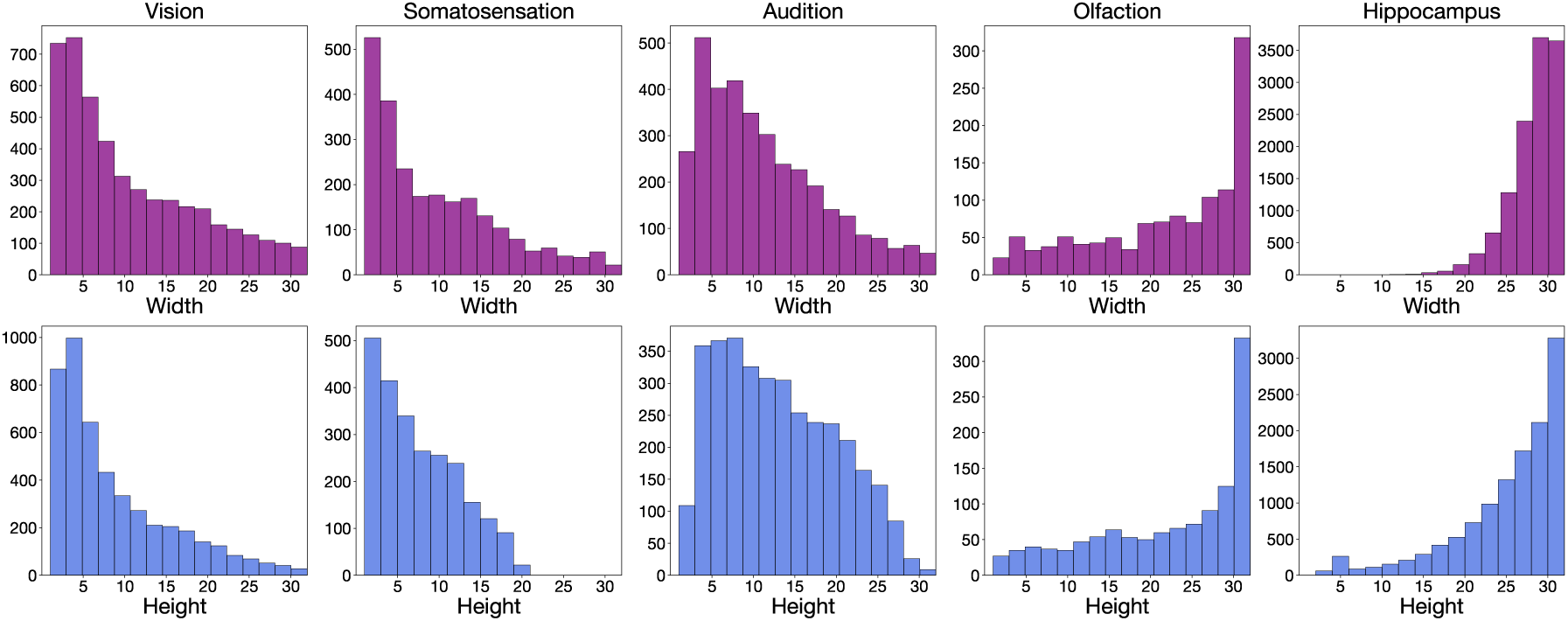
Distribution of receptive field sizes (height and weight shown separately) of network units after training a fully-connected network for each of the five modalities. Receptive field sizes are defined as the minimal bounding box surrounding all non-zero values within a given row of the layer’s weight matrix arranged in two dimensions. Note the difference between the shape of the distributions in the spatiotopic modalities and olfaction/hippocampus

### Hyperparameter sweep of the self-organizing map algorithm

The configurable parameters in the self-organizing map algorithm (Algorithm 1 in supplementary methods) are the initial learning rate and the map size. All other parameters of the algorithm are computed as a function of these two. The results presented in the main text use an initial learning rate of 0.1 and a map size of 25 × 25. The results are consistent across variations of these settings. A parameter sweep is presented below.

Figure S3 shows the indices for the spatiotopic order of the maps and the locality of their receptive fields for the five modalities across different values of the initial learning rate. The same patterns as in Figure 5f-g of the main text are seen across these settings.

**Figure S3:**
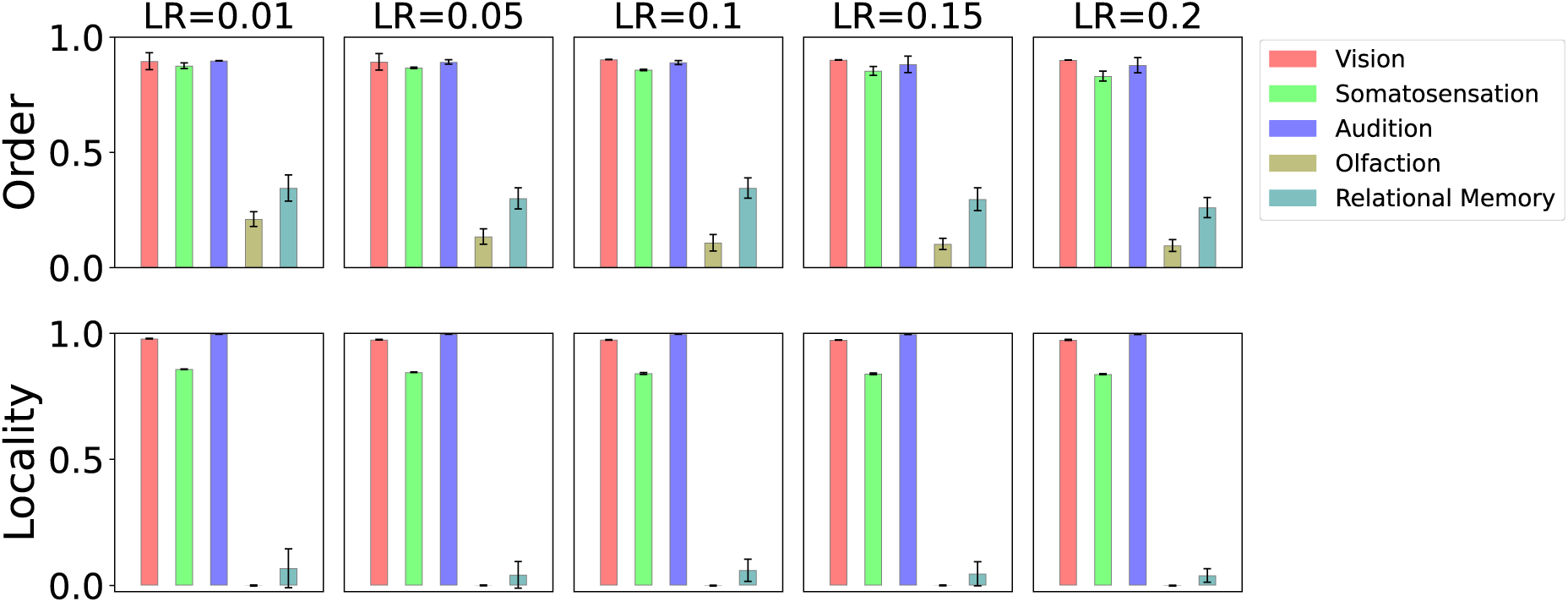
Spatiotopic order of the maps and the locality of their receptive fields in the five modalities across different values of the learning rate (LR). Plotted are the means and standard deviations across ten simulations with different random seeds.

Figure S4 shows that different map sizes produce the same overall results.

**Figure S4:**
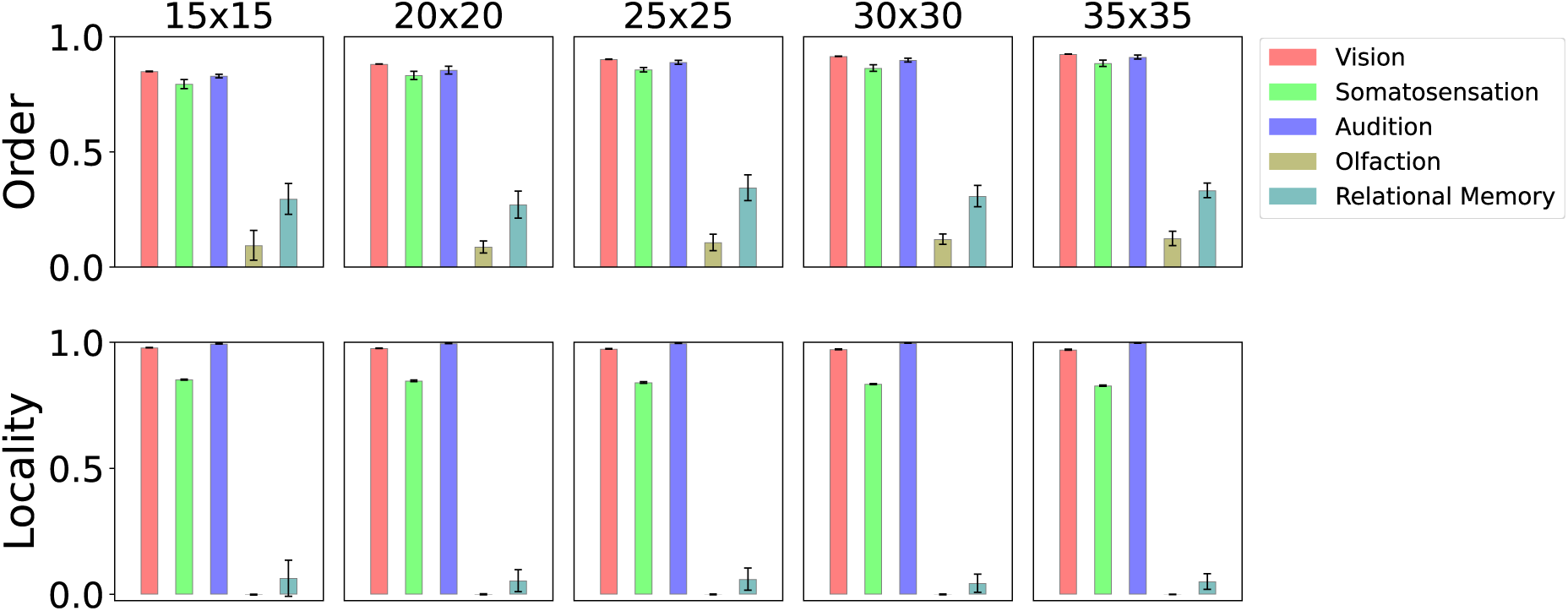
Spatiotopic order of the maps and the locality of their receptive fields in the five modalities across different map sizes. Plotted are the means and standard deviations across ten simulations with different random seeds.

The same overall results are also observed for different patch sizes (used to probe the organization of the map) as well as for PCA map initialization, where the initial weights of the map units are set to the first principal components of the map input instead of random values.

### Hyperparameter sweep of the evolutionary algorithm

The configurable parameters in the evolutionary algorithm are (1) the fraction of best-performing networks in a given generation that persist to the next generation, and (2) the extent of the variation of the boundary location around that of the highest-performing network. In the following, these two parameters are referred to as *k* and *σ* respectively. In Figure 6 of the main text, values of *k* = 0.5 and *σ* = 0.1 are used; *σ* = 0.1 means that new networks are formed by sampling uniformly within ±0.1 of the fittest network’s boundary location, which lies in the range [0, 1], 0 corresponding to one end of the network and 1 corresponding to the other end. The results are consistent across variations of these parameters. A parameter sweep is presented below.

Figure S5 sweeps through different values of *k*. The pattern of covariation between network components stays the same across settings. Figure S6 sweeps through different values of *σ*, again showing the same trends.

When generating new networks, the total number of parameters allocated to the olfactory network is allowed to drift randomly by up to a percentage *δ*, resulting in an additional degree of variability among the new networks. In Figure 6 of the main text, a value of *δ* = 10% is used. The same overall trend is observed for different settings of *δ*, as shown in Figure S7.

**Figure S5:**
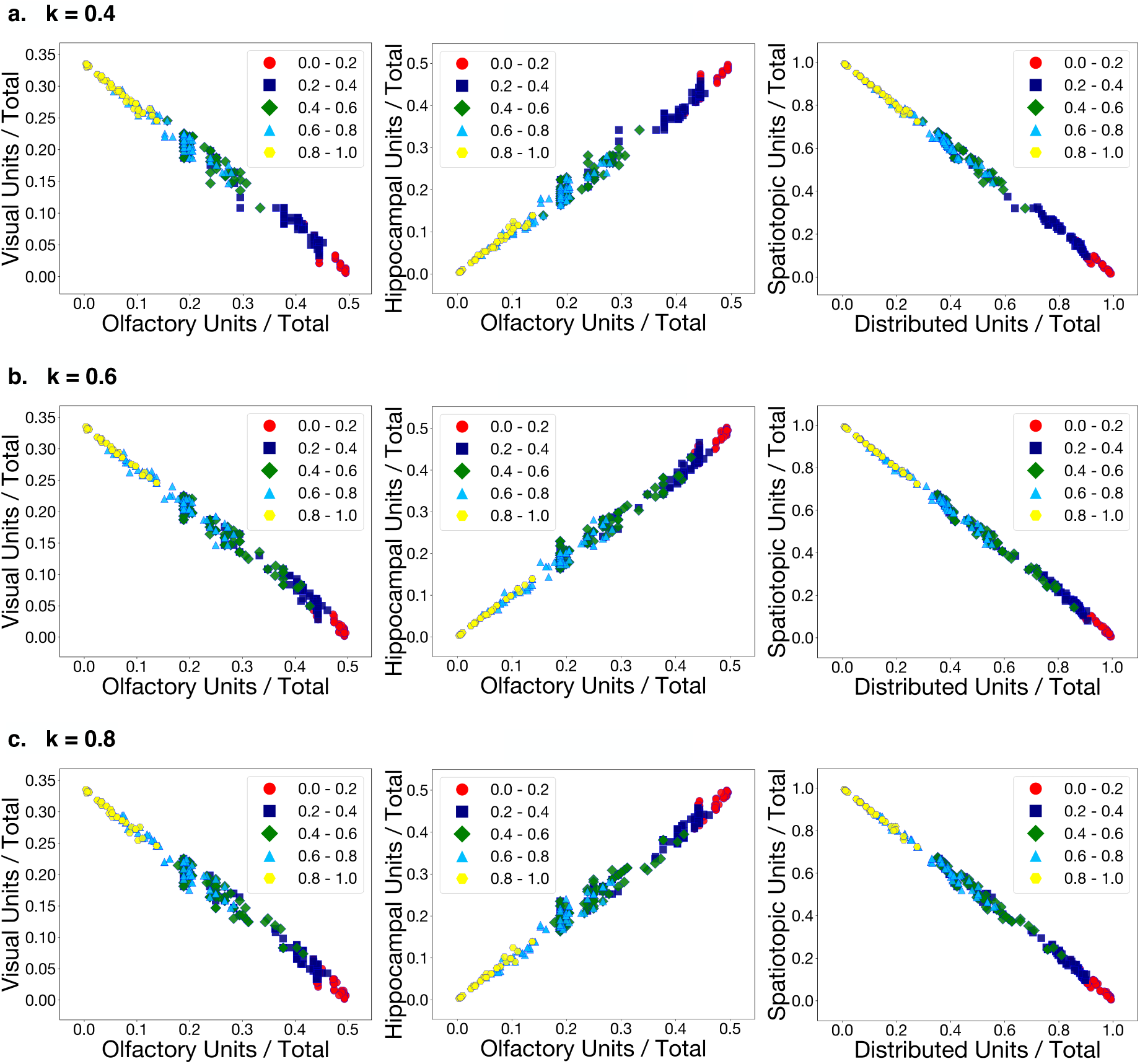
Covariation patterns resulting from the evolutionary algorithm for different values of the parameter *k*, which determines the fraction of best performing networks in a given generation that persist to the next generation.

**Figure S6:**
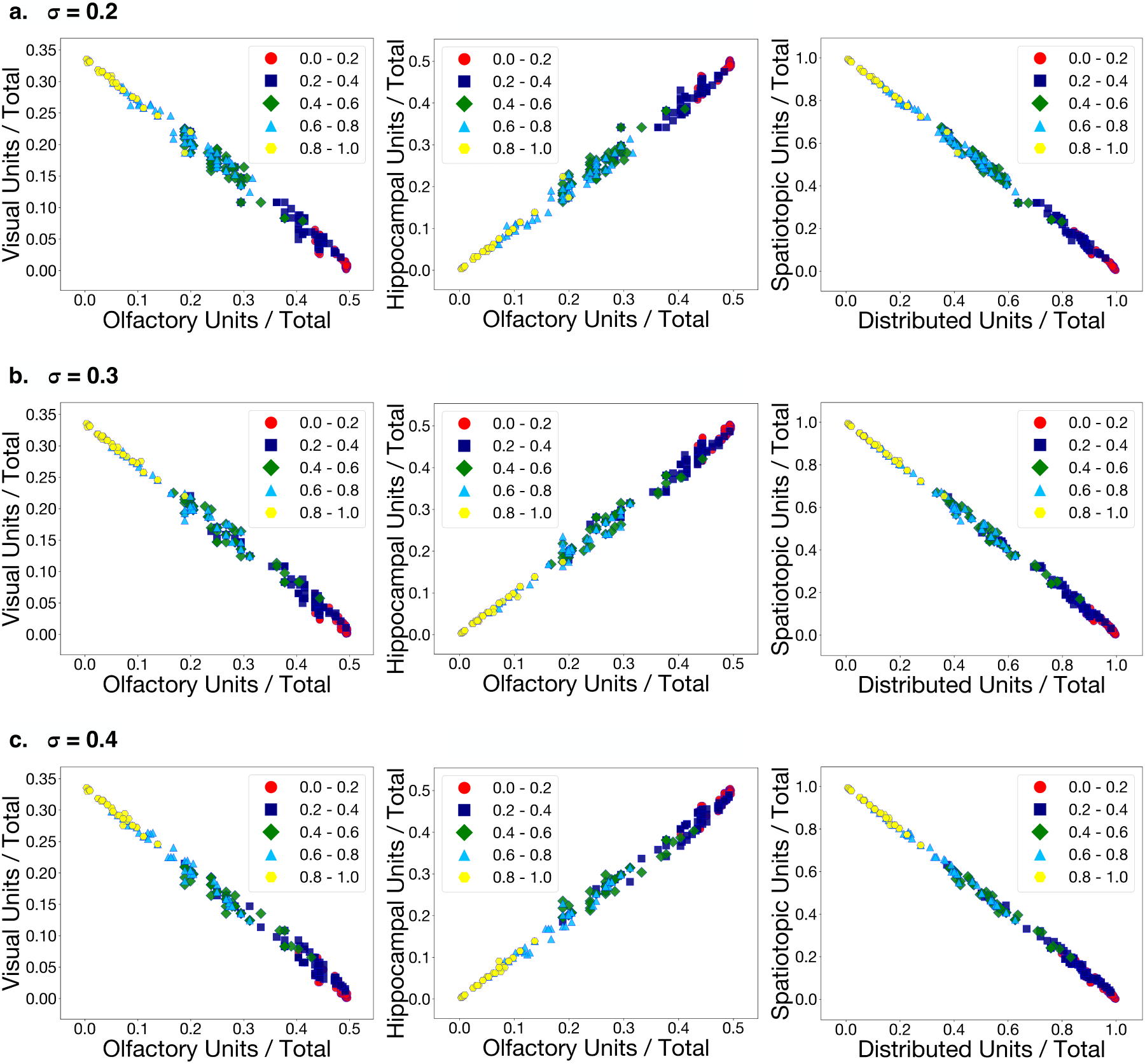
Covariation patterns resulting from the evolutionary algorithm for different values of the parameter *σ*, which determines the extent of the variation of the boundary location around that of the highestperforming network.

**Figure S7:**
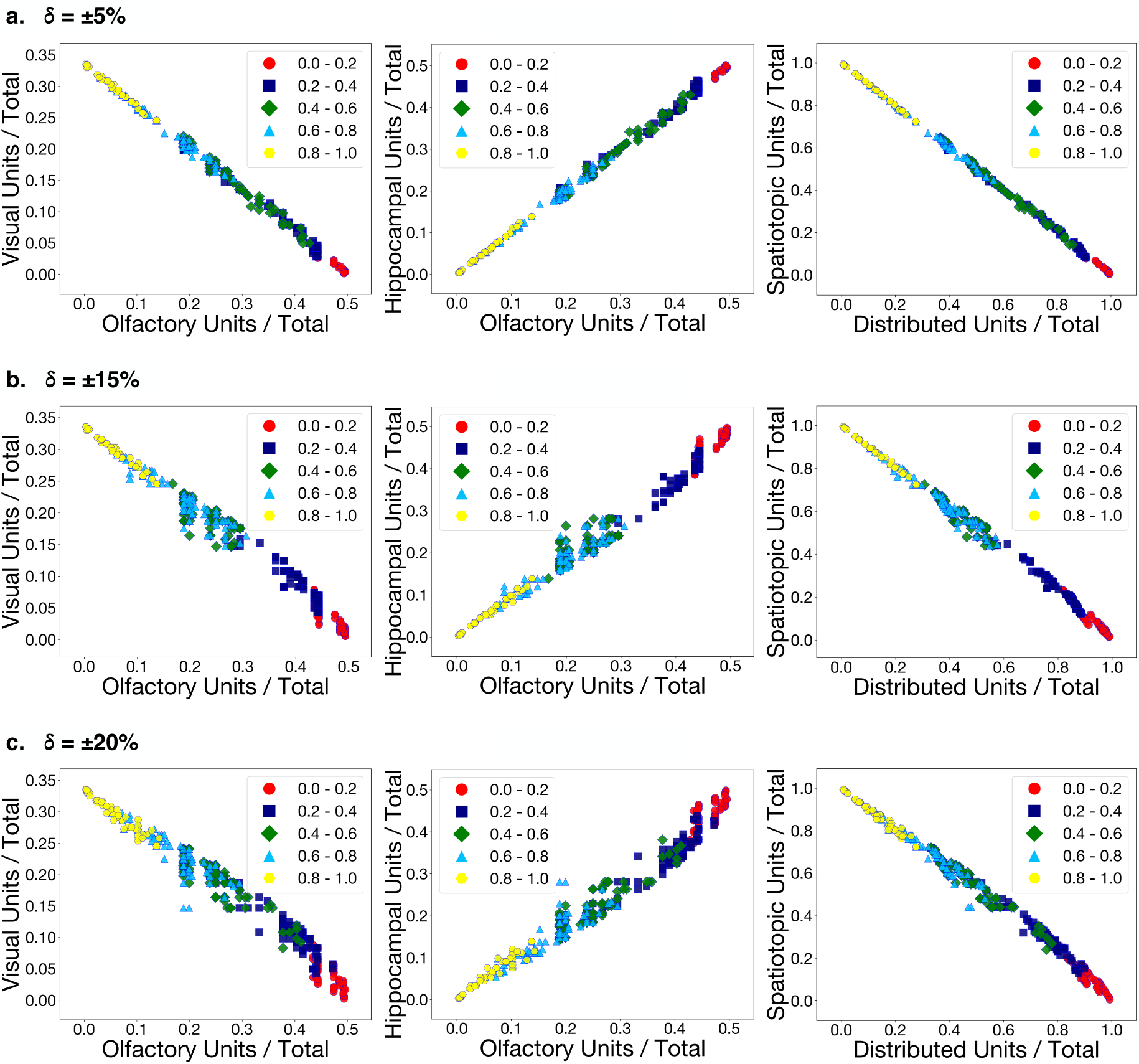
Covariation patterns resulting from the evolutionary algorithm for different values of the drift parameter *δ*.

